# Processive movement of Actin by Biased Polymerization: A new paradigm of Axonal Transport

**DOI:** 10.1101/212449

**Authors:** Nilaj Chakrabarty, Pankaj Dubey, Yong Tang, Archan Ganguly, Kelsey Ladt, Christophe Leterrier, Peter Jung, Subhojit Roy

## Abstract

Classic pulse-chase studies have shown that actin is conveyed in slow axonal transport, but the mechanistic basis for this movement is unknown. Recently, we reported that axonal actin was surprisingly dynamic, with focal assembly/dis-assembly events (“hotspots”) and elongating polymers along the axon-shaft (“trails”). Using a combination of live imaging, super-resolution microscopy, and modeling, here we explore how these axonal actin dynamics can lead to processive transport. We found abundant actin nucleation, along with a slow, anterogradely-biased flow of actin in axon-shafts. Starting with first principles of monomer/filament assembly – and incorporating imaging data – we generated a quantitative model simulating axonal hotspots and trails. Our simulations predict that the axonal actin dynamics indeed lead to an anterogradely-biased flow of the population, at rates consistent with slow transport. Collectively, the data point to a surprising scenario where local assembly and biased polymerization generate the slow axonal transport of actin. This mechanism is distinct from polymer-sliding, and seems well suited to convey highly dynamic cytoskeletal cargoes.

**Acknowledgements:** **This work was supported by an NIH grant to SR** (R01NS075233). The authors thank Stephanie Gupton (UNC) for the Mena/Vasp constructs.

## INTRODUCTION

Actin is a key cytoskeletal protein in neurons, with important roles in axon growth and synaptic homeostasis (Coles and Bradke, 2015; Gomez and Letourneau, 2014; Kevenaar and Hoogenraad, 2015). Although small amounts of actin can be synthesized locally in response to axonal guidance cues (Jung et al., 2014), the vast majority of actin – along with the other major cytoskeletal proteins tubulin and neurofilaments – is made in the neuronal soma and conveyed into axons via slow axonal transport, as shown by classic in vivo pulse-chase radiolabeling studies in many different organisms (Black and Lasek, 1979; McQuarrie et al., 1986; Oblinger, 1988; Tashiro and Komiya, 1992); reviewed in Galbraith and Gallant, 2000; Roy, 2013. Although these studies defined the overall transport of actin, underlying mechanisms remained obscure, as radioisotopic labeling cannot visualize cargo-movement.

More recently, live imaging with fluorescent-tagged probes have begun to reveal the mechanistic basis of cytoskeletal slow axonal transport. Imaging of GFP-tagged neurofilaments revealed that neurofilament polymers move rapidly but intermittently in axons, resulting in a slow overall movement of the population (Roy et al., 2000; Wang et al., 2000) – the “Stop and Go” model, (Brown, 2000). Similarly, short, motile structures resembling microtubules also move intermittently in axons (He et al., 2005; Wang and Brown, 2002). Conceptually, these imaging studies advocate a mechanism where cytoskeletal polymers assemble in the neuronal soma, and the assembled polymers are translocated into axons by motor proteins. Unfortunately, such straightforward imaging strategies have not been useful for deciphering actin transport. One issue is that actin is much more dynamic than neurofilaments and neuronal microtubules, and a significant fraction (about half) exists as monomers (Morris and Lasek, 1984). Consequently, GFP-tagging of monomeric actin typically reveals a diffuse glow in the axon, with few discernible structures (Okabe and Hirokawa, 1990). A further confounding factor is that GFP-tagged actin may not report all actin behaviors – particularly ones mediated by formins (Chen et al., 2012).

Using probes that selectively bind to filamentous actin, we recently visualized actin dynamics in axons of cultured hippocampal neurons (Ganguly et al., 2015). We found that actin continuously polymerizes and de-polymerizes at micron-sized "hotspots" along the axon, spaced ~ 3-4 µm apart. Interestingly, the hotspots were colocalized with stationary axonal endosomes, suggesting nucleation of actin on the surface of vesicles. In addition, we saw rapidly elongating actin polymers extending along the axon-shaft ("actin trails"). Actin trails were formin (but not Arp 2/3) dependent; typically originated from the hotspots; and helped enrich actin at presynaptic boutons. Based on these data, we proposed a model where axonal actin nucleates on the surface of stationary endosomes, providing the nidus for polymers elongating along the axon shaft.

Though axon shafts have other stable actin structures such as “actin rings” (Xu et al. 2013; Ganguly et al., 2015), the abovementioned hotspots/trails are the only known dynamic elements in mature axons. Thus it seems reasonable to imagine that the network of hotspots and trails would somehow lead to the axonal transport of actin. However, this is not straightforward to conceptualize, as actin dynamics in axons are not like neurofilaments and microtubules, where stable polymers are simply translocated by motors. Instead, the nature of actin assembly/disassembly – occurring on the timescale of seconds – requires one to consider the biophysics of this exchange. Using a combination of imaging and quantitative modeling, here we show that a dynamic but polarized assembly of actin in axons can indeed lead to a slow anterograde bias of the population, at rates consistent with slow transport. Our data point to an unconventional axonal transport paradigm that is fundamentally based on dynamic assembly, and yet results in biased transit of the population.

## RESULTS AND DISCUSSION

### Dynamics of local actin assembly and polymer-elongation in axon shafts

Kymographs in **figure 1A** show examples of axons transfected with GFP:Utr-CH (GFP bound to the calponin homology domain of utrophin), a probe that selectively labels actin filaments (Burkel et al., 2007). Note two key features: 1) Repeated assembly/dis-assembly of actin in discrete microscopic zones along the length of the axon, appearing as vertical interrupted lines in the kymographs (hotspots); and 2) Bidirectionally elongating actin polymers, appearing as diagonal ‘plumes’ in the kymographs (actin trails). Also note that the actin trails often originate from hotspots (some marked with red dashed circles in **fig. 1A**; also see (Ganguly et al., 2015). Actin hotspots/trails are seen with other actin probes such as Lifeact (Ganguly et al., 2015); as well as in *C. elegans* axons in vivo (Sood et al., 2017). Interestingly, though the average elongation rate of actin trails was similar in both directions, the frequency of anterogradely elongating actin filaments was slightly higher (~ 55% elongated anterogradely, **fig. 1B**).

**Figure 1:**
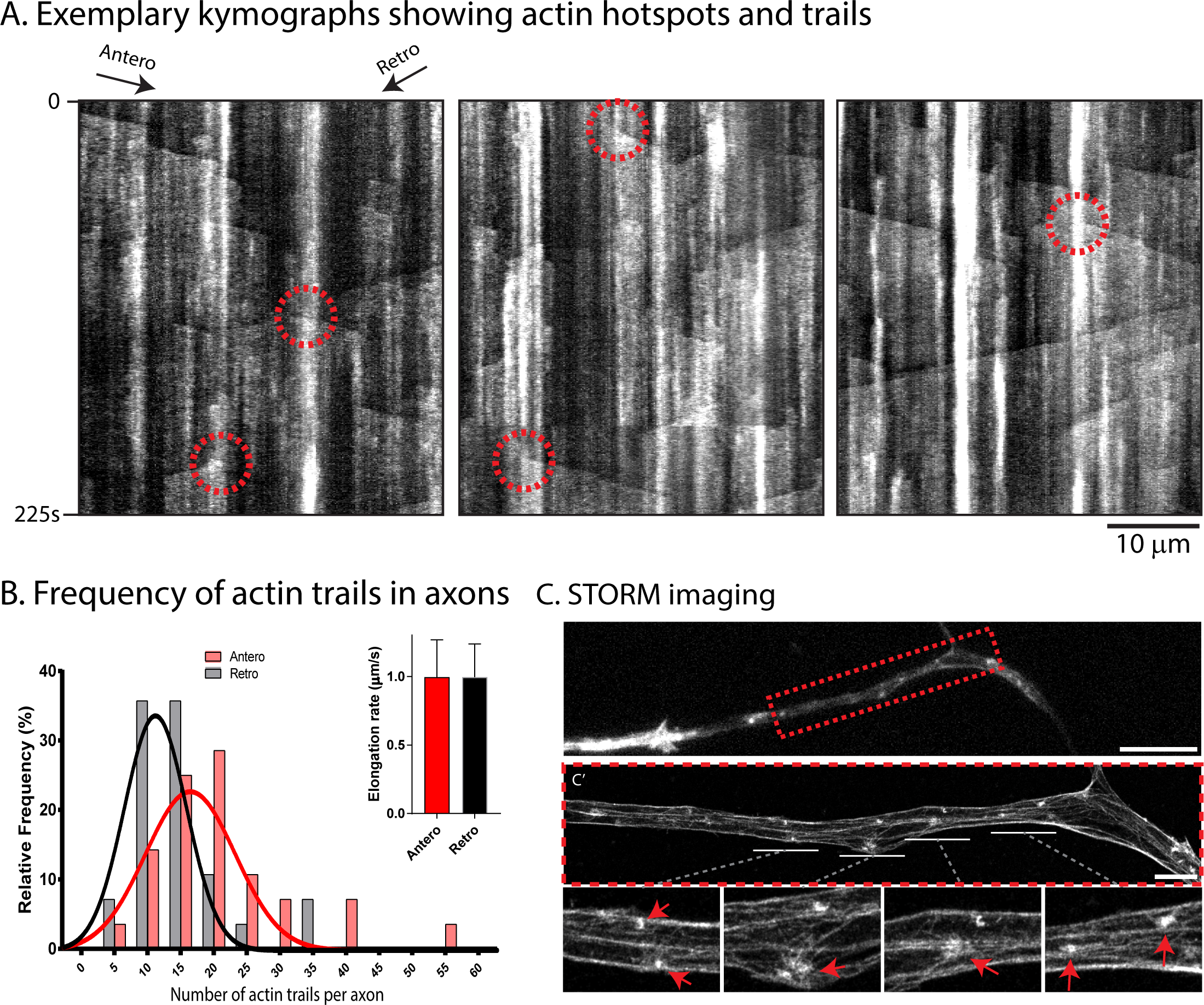
Dynamics and organization of actin in axon shafts. **(A)** Kymographs from axons transfected with GFP:Utr-CH to label filamentous actin. Distance is along the x-axis, and elapsed time is shown on y-axis. Note vertical “on/off” events representing dynamic local assembly (actin hotspots); and diagonal plumes of fluorescence representing bidirectionally elongating actin polymers (actin trails). Also note that actin trails often emerge from where the hotspots are situated (some marked by dashed circles; also see Ganguly et al., 2015). **(B)** Anterograde and retrograde frequency of actin trails (histogram) and polymer elongation rate (graph, upper right). Note the slightly higher frequency of anterograde trails (58/42%; mean elongation rates were 0.99 +/-0.001 µm/s in both directions; data from ~ 1000 total events, adapted from Ganguly et al., 2015). **(C)** Axon from a DIV-3 neuron labeled with phalloidin (C’ shows STORM image of region within dashed red box). Note that several clusters of actin with radiating filaments are seen in the STORM images (arrows in zoomed insets below); likely corresponding to actin hotspots seen by live imaging. Scale bar: C, top = 10 um; C′ = 2 um.

An important concept emerging from our experiments is that axon shafts have microscopic zones where actin is nucleated. In previous studies using stochastic optical reconstruction microscopy (STORM) to examine actin in axons, we found a two-tier distribution – circumferential actin rings underneath the plasma membrane, and a deeper network of linear actin filaments (Ganguly et al., 2015). Examining the latter in younger axons that have relatively thicker profiles, we saw discrete clusters of actin along the axon-length, with “aster-like” radiating actin filaments (**fig. 1C**). The spatial distribution and morphology of these clusters in the STORM images strongly suggest that they represent the ‘hotspots’ that we see by live imaging, supporting the idea that axons have discrete foci where actin is nucleated. Similar axonal actin clusters are seen in more mature axons as well (**Supp. fig. 1**). Interestingly, the STORM data indicate that many of these actin clusters are close to the plasma membrane – a feature that cannot be appreciated by diffraction-limited microscopy – suggesting that there may be anatomical and/or mechanistic links between actin rings and actin trails.

Next we directly examined sites of new actin incorporation in axons. Though previous studies have examined sites of actin nucleation in axons, they have exclusively focused on the growth cone, showing that new actin barbed-ends are incorporated to the leading-edge of growth cones (Marsick et al., 2010). To determine if such nucleation can also occur along the axon shaft, we used an established method to highlight newly incorporated actin barbed ends in cells (Marsick and Letourneau, 2011; Symons and Mitchison, 1991). Briefly, in this technique, cells are incubated with rhodamine-labeled actin monomers and a mild detergent (to allow the entry of labeled monomers into cells), and then fixed and imaged (see schematic in **fig. 2A** and “methods”). Consistent with previous studies, we saw preferential incorporation of labeled monomers along filopodial tips in CAD cells (**fig. 2B**). To ensure that our experiments reported local incorporation of actin monomers along axon shafts, we adapted the barbed-end labeling technique in microfluidic chambers where axon shafts can be physically and fluidically isolated from somato-dendritic domains. The rhodamine-labeled actin monomers are only added to the axonal chamber in these experiments (see **fig. 2C**); thus any labeling in axons is due to local monomer-assembly (and not transport/diffusion from the soma/dendrites). Indeed, there was extensive rhodamine-actin labeling in axons (**fig. 2C**). Interestingly, both punctate and elongated actin structures were seen, likely because the rhodamine actin also labeled actin trails that elongated during the two-minute incubation period with the labeled monomers (arrowheads in zoomed inset, **fig. 2C**). Furthermore, the barbed-end binding proteins Ena/Vasp (Breitsprecher et al., 2011) were precisely localized to the hotspots (**fig. 2D**), suggesting that actin trails elongate bidirectionally with their barbed-ends facing the hotspots (see schematic in **fig. 2E**).

**Figure 2:**
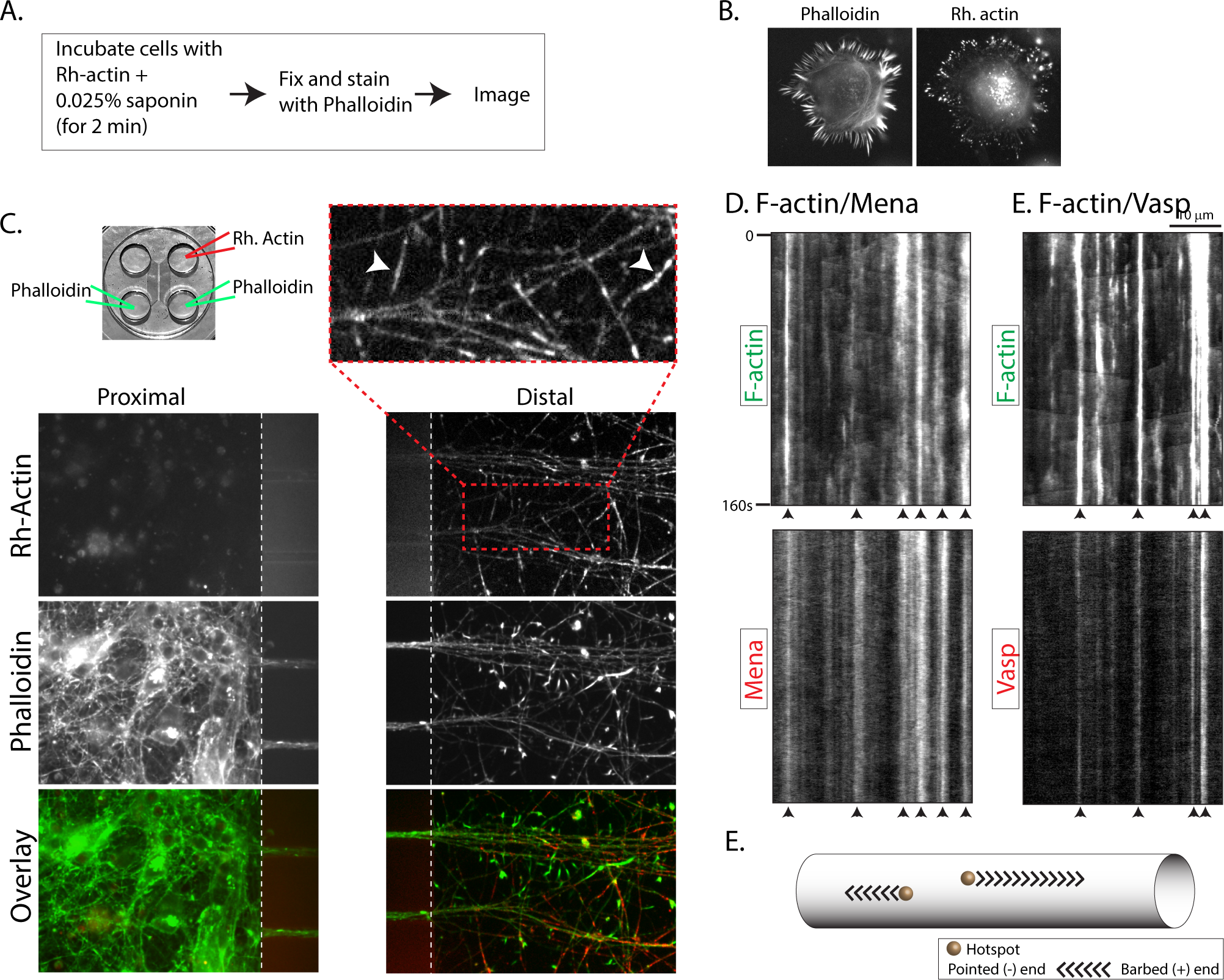
Labeling of newly incorporated barbed-ends in axon shafts. **(A)** Protocol to label newly incorporated barbed-ends of actin in cells. **(B)** Representative images of CAD cells after barbed-end labeling. Note that the newly incorporated barbed ends (labeled with rhodamine-actin) are enriched at filopodial tips. **(C)** <u>Top left</u>: Microfluidic device to separate axons from somatodendritic compartments. To specifically label barbed ends in axons, rhodamine-actin was loaded only into the axonal chamber, and phalloidin staining was performed on both chambers, as described in “methods”. <u>Bottom</u>: Representative images showing rhodamine-actin and phalloidin labeling in somatodendritic (left panels) and axonal chambers (right panels). Note numerous small rhodamine-actin puncta in axons, indicating local incorporation of monomers (zoomed image on top right). Also note some elongated structures (arrowheads in zoomed image), likely representing rhodamine label incorporated into elongating actin filaments. **(D)** Kymographs from axons co-transfected with GFP:Utr-CH and mCherry:Mena (or mCherry:Vasp), and imaged near-simultaneously as described in “methods”. Note striking co-localization of actin hotspots with Mena/Vasp (some marked with arrowheads). **(E)** Schematic of trail elongation; note that trails grow with their barbed-ends facing the hotspots.

### Biased anterograde flow of axonal actin

Although radiolabeling studies defined the various classes of axonal transport, the movement cannot be visualized by these methods. More recently, we developed a quantitative imaging assay to visualize slow axonal transport in cultured neurons (Ganguly et al., 2017; Scott et al., 2011; Tang et al. 2013). In these experiments, proteins known to move in slow axonal transport (as shown by radiolabeling techniques) are tagged to photoactivatable GFP (PAGFP) and transfected in cultured neurons. Thereafter, a discrete pool of molecules is photoactivated in the axon, and the resultant fluorescent pool is tracked by live imaging. Dispersion of fluorescence is biased towards the axon-tip, which can be quantified by analyzing the anterograde shift in the center of fluorescence-intensity over time (“intensity-center shift”). The biased movement is energy dependent, and distinct from the rapid and unbiased diffusion of PAGFP alone (Scott et al. 2011). To determine if there is a similar bias of actin in axons, we transfected hippocampal neurons with PAGFP:Utr-CH (and soluble mCherry as a volume marker), and performed the photoactivation experiments (**fig. 3A**). Indeed, there was a slow, anterograde bias of the labeled actin population, as shown in **figure 3B-D**. The overall rate of actin egress, as calculated from the slope of the intensity-center shift, was ~ 0.34 mm/day, in line with previous pulse-chase radiolabeling studies in CNS neurons [~ 0.4 mm/day, see (Oblinger, 1988)]. Photobleaching of GFP:Utr-CH in axons also suggested a biased flow of actin in axons (**fig. 3E**), though this was difficult to quantify.

**Figure 3:**
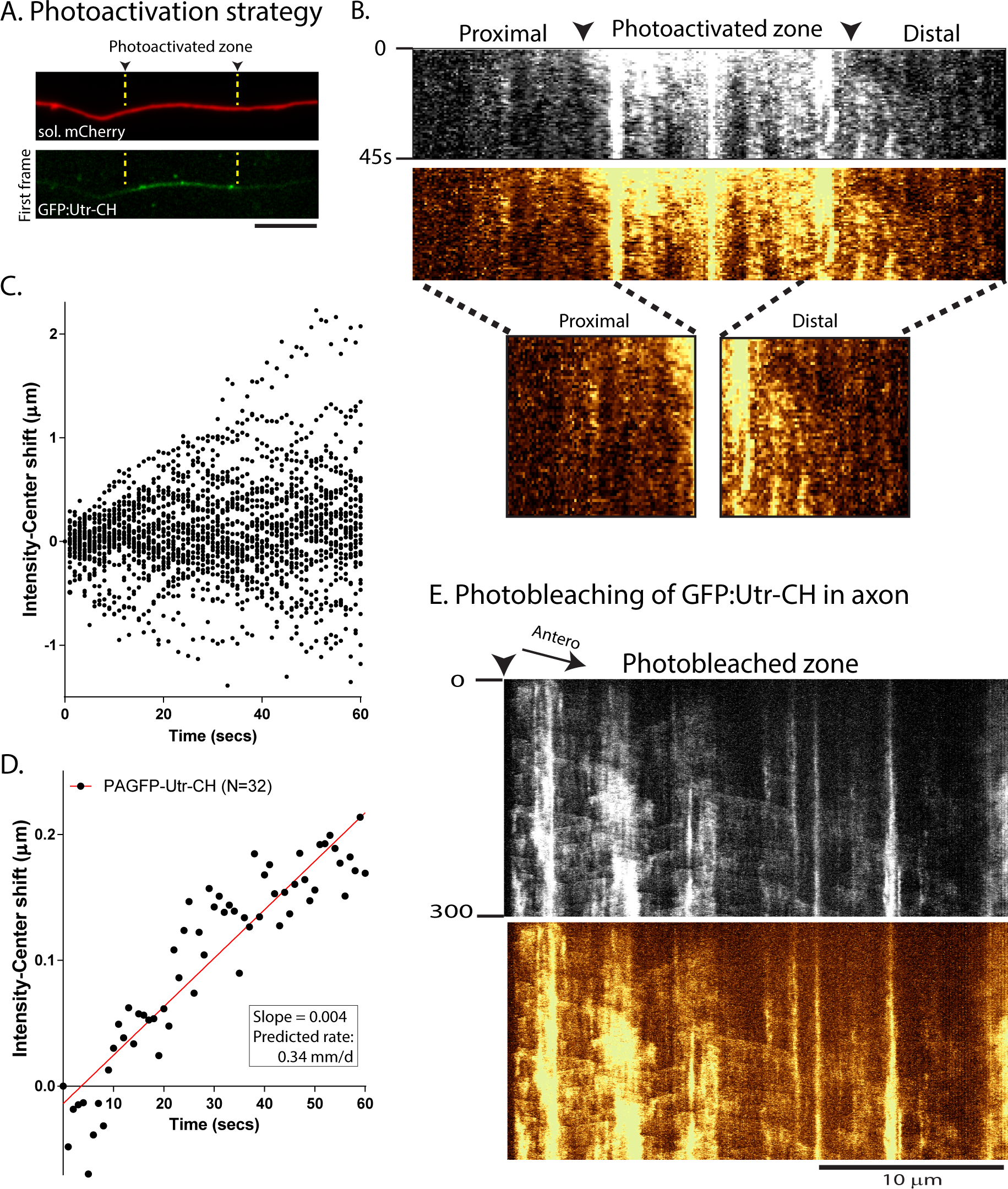
Anterograde bias of actin in axons. **(A)** Representative images of an axon co-transfected with PAGFP:Utr-CH and soluble mCherry. Note that a discrete region of interest is photoactivated (green), and the mobility of the fluorescent pool is analyzed over time. **(B)** Greyscale (top) and pseudo-colored (bottom) kymographs from a GFP:Utr-CH photoactivation experiment. The photoactivated zone is marked by arrowheads, and elapsed time in seconds is shown on left. Zoomed insets (far bottom) highlight the anterogradely-biased dispersion of axonal actin. **(C-D)** Quantification of photoactivation experiments by analyzing the displacement in the centroid of fluorescence over time (“intensity center shift”, see methods for more details). Raw (C) and mean (D) intensity center shifts reveal a slow, anterograde bias of the actin population. **(E)** Kymograph of an axon showing photobleaching and recovery of bleached zone in a neuron transfected with GFP:Utr-CH. Note the anterogradely-biased flow of fluorescence; a few actin trails are also visible.

### Simulation of axonal actin hotspots and trails

The mechanistic model emerging from our preceding experiments is: 1) Axons have discrete foci where actin monomers are nucleated (hotspots); 2) Actin filaments elongate bi-directionally from the hotspots, with barbed ends facing the hotspots; and 3) Actin has an overall, slow anterograde bias in axons. *How does the dynamic actin network of hotspots and trails lead to the slow, biased flow of the population?* Mechanistically, the underlying process is likely complex, involving actin monomer/polymer exchange, assembly/dis-assembly of hotspots/trails, and elongation of actin filaments. To address this, we first designed a robust simulation of actin hotspots and trails in axons, and then performed “virtual photoactivation experiments” – asking if hotspots/trails could lead to an overall biased axonal transport. The simulations employed established biophysical principles of actin monomer/filament assembly; incorporating parameters from our imaging data. Specifically, multiple virtual actin hotspots were allowed to originate linearly along a hypothetical axon-cylinder (axon-thickness and distance between the nucleating zones was based on imaging data from (Ganguly et al., 2015); with polymers extending from these hotspots. A schematic of the modeling is shown in **fig. 4A**, left). Note that monomers nucleate with their barbed-ends facing the hotspots, with polymers extending in both anterograde and retrograde directions. Also note that in this scenario, addition of new monomers at the barbed ends of the elongating trails will lead to translocation of individual monomers towards the pointed end of the growing filament (**fig. 4A** – left, dashed inset at bottom; also see Supp. Movie 1).

**Fig 4:**
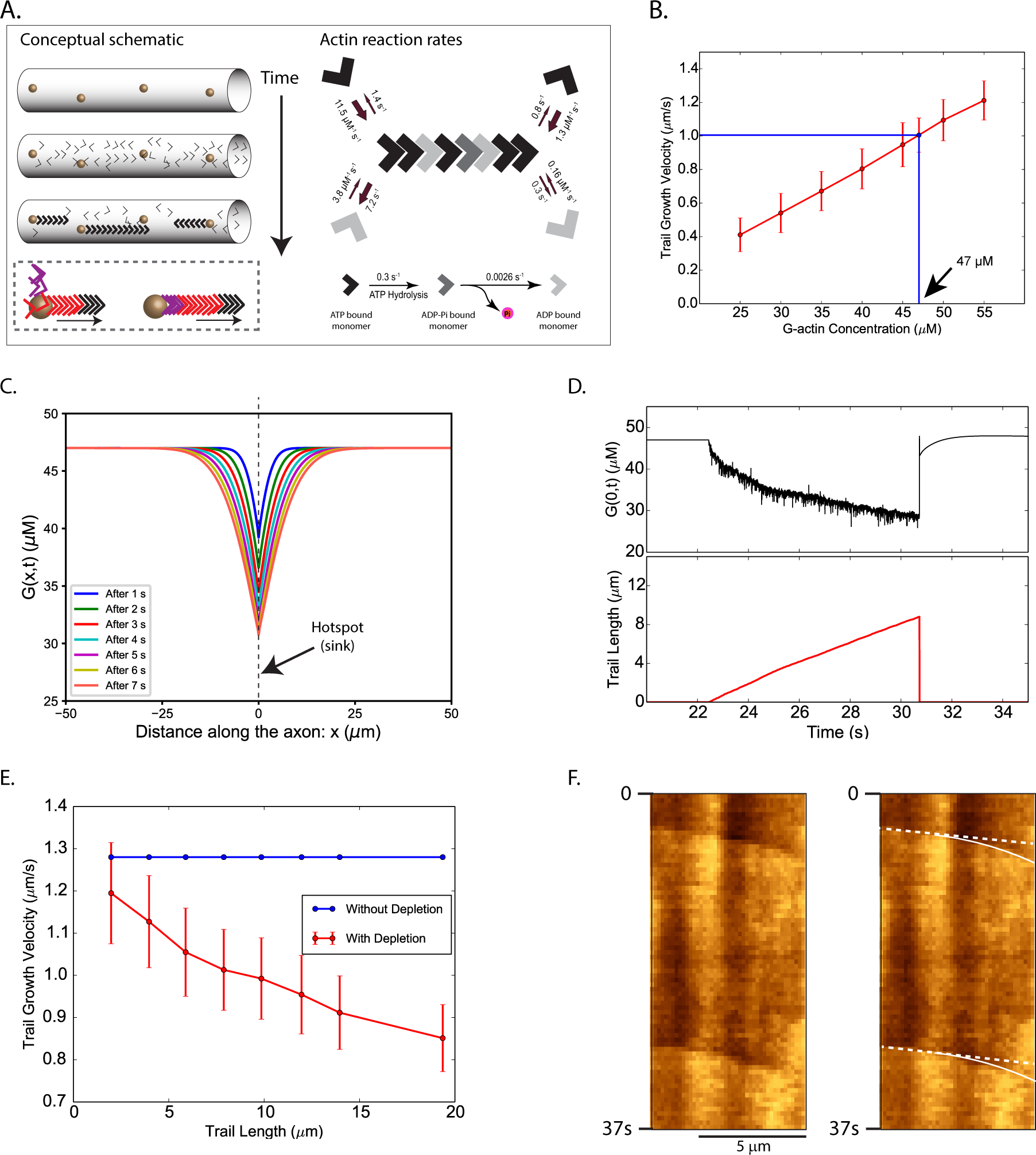
Modeling of axonal actin hotspots and trails. **(A)** <u>Left</u>: Conceptual schematic of our modeling. Stationary hotspots (y ell ow sph eres) are locali zed unif ormly along the axon; and actin trail s are nucleated on the hotspots, with barbed ends facing the hotspots. Note that as new monomers (colored arrowheads) are incorporated at the barbed end, there is progressive translocation of indi vid ual monomers towards the pointed end (box below). <u>Right</u>: Known assembly, disassembly, and summary of reaction rates of actin. **(B)** Determination of basal actin monomer concentration in the axon. Average elongation velocity of actin trail s increases li nearly (red li ne) with the basal monomer concentration. At a basal concentration of 47 µM the average elongation velocity matches the experimentally observed average trail elongation velocity of ~ 1 µm/s (blue line). **(C)** Actin monomer concentration profiles along the axon during the growth of an actin trail. Multiple colored lines show declining concentration of actin monomers along the axon shaft at successive time-points after trail-nucleation. Note that the hotspot is located at x = 0 and the trail grows with its barbed end attached to this location. The trail collapses about 9 seconds after nucleation, growing to a length of 8.8 µm. This wide concentration-gradient drives in monomer from regions of higher concentration towards the hotspot. **(D)** Actin monomer depletion during trail elongation. An actin trail is randomly nucleated at t = 22.4 s and the trail grows (lower panel, red trace), monomer depletion from the barbed end (upper panel, black trace) slows down the rate of trail elongation. Note that fluctuations of actin-monomer concentration reflect monomers randomly attaching/detaching from the barbed end. As the trail collapses at t = 30.7 s, actin-monomer concentration spikes back up. **(E)** Effect of actin-monomer depletion on trail elongation (basal 47 µM monomer concentration). Note that trail elongation velocities decrease with increasing trail-lengths (red line). **(F)** A kymograph of GFP:Utr-CH in an axon, depicting slowing of trails over time (note change in slope, marked by curved line). Distance is on x-axis and time is on the y-axis.

The on/off kinetics of actin in our model is in accordance with known biophysical properties of monomer/polymer exchange. Briefly, actin monomers can bind to either ATP or ADP in its cleft, with distinct association and dissociation rates from the ends of actin polymers (Pollard, 1986). In the filaments, ATP-actin can hydrolyze irreversibly to form ADP-Pi-actin (Carlier et al., 1988) which undergoes Pi release to form ADP-actin (Carlier and Pantaloni, 1986). The known association, dissociation, and hydrolysis rate constants of ATP-actin and ADP-actin are summarized in **figure 4A**, right (see (Blanchoin and Pollard, 2002; Carlier and Pantaloni, 1986; Pollard, 1986). The ATP/ADP ratio in neurons is very high – thought to sustain its extreme energy requirements (Tantama et al., 2013) – and we assume that ATP-bound monomers are the only species that can be added to elongating axonal filaments. Once bound to the filament, subunits in our model randomly undergo hydrolysis at a rate of 0.3/*s*, and release the phosphate group at a rate of 0.0026/*s* (Carlier and Pantaloni, 1986). It is known that actin filaments can linearly grow up-to 17.7 µm before they start bending (called “persistence length”, see (Gittes et al., 1993). Since the lengths of the axonal actin trails is much smaller (average 8.8 µm, see (Ganguly et al., 2015), we modeled them as one-dimensional (1-D) linear filaments elongating along the axon shaft (also note that they appear linear by live imaging, see **fig. 1A** for example). Given that the typical diameter, *d*, of an axon visualized in our imaging experiment is approximately 200 *nm*, the maximum angle of an elongating actin trail (of length *L*_*t*_) with the long-axis of the axon is only ~ 1 ° [*θ* = *tan*^−1^(*d*/*L*_*t*_) = 1.2° for a typical trail length of 8.8 *μm*], justifying a 1-D mathematical model of axonal actin trails.

The nucleation of actin trails, and their subsequent elongation by incorporating monomers from the axonal pool was modeled by Markov processes. Nucleation rates of anterograde and retrograde trails were calculated from imaging studies of GFP:Utr-CH (Ganguly et al., 2015). Specifically, the average nucleation rate (*r*_*n*_) of anterograde and retrograde trails (for a given hotspot) was *r*_*n,a*_ = 0.001885/*s* and *r*_*n,r*_ = 0.001381/*s* respectively (see **Supplementary Materials and Methods eq. 1**). Probabilities of trail nucleation, and of competing association and dissociation reactions were calculated at each time-step using the nucleation rates *r*_*n,a*_ and *r*_*n,r*_ and established reaction-rates. The effect of incorporating actin monomers from the axonal pool into the elongating trails, and the subsequent release of actin back into the axonal pool was modeled using a 1-D diffusion equation with sinks and sources. Based on previous studies, the diffusion coefficient of actin monomers was assumed to be 6 *μm*^2^/*s* (McGrath et al., 1998) Since the elongation rate of actin trails in axons is known [~ 1 µm/s, see **fig. 1** and (Ganguly et al., 2015)], we estimate that a monomer concentration of 47 *µ*M would be needed to sustain the polymerization at this rate (**fig. 4B**). Note that though the concentration of actin in cultured hippocampal axons is unknown, our estimate is within the range of actin concentration in other cell types (for example, concentration of monomeric actin in chick embryonic neurons is reported to be 30-37 µM, see (Devineni et al., 1999). For more details of the simulation, **see Supplementary Materials and Methods** and **Supplementary Figure 2**.

In our imaging experiments, actin trails invariably collapse after elongating for a few microns [see **fig. 1A** and (Ganguly et al., 2015)]. In our simulations however, if unchecked, the trails would probably grow indefinitely, indicating that the collapse is not a simple consequence of monomer depletion, but is mediated by yet unknown mechanisms. To more closely mimic the actual experimental data, we forced the simulated elongating trails to collapse after reaching predetermined lengths (determined from the distribution-range of “polymer length” data from (Ganguly et al., 2015), average 8.8 µm). **Figures 4C, D** highlight the kinetics monomer/polymer exchange in our model when an actin trail elongates and collapses. Note that as the filament grows, the monomer pool at the hotspot is depleted, supplying subunits to the elongating trail (**fig. 4C**). The dynamic actin subunit/polymer in our model is shown in **figure 4D**. Here, note that the actin monomer pool (black trace) is depleted as the filament grows over a few seconds (red trace). Also note that though monomer-depletion cannot explain the abrupt collapse of trails in our axons, our modeling suggests that the rate of polymer elongation should slow down as the filament grows (**fig. 4D, E**). Indeed upon closely examining our GFP:Utr-CH kymographs we saw examples where the elongating actin trails gradually slowed down (**fig. 4F**). For molecular simulations of the monomer/polymer exchange, and elongation/collapse of actin trails, see **Supplementary Movies 1 and 2**.

### Simulation of axonal actin photoactivation experiments

To determine if the axonal actin dynamics (hotspots/trails) can generate an overall biased egress of the population, we performed virtual photoactivation experiments in axons with simulated hotspots and trails, and asked if there was a shift in the resultant “virtual intensity center shift” (as in our actual imaging experiments, see **fig. 3**). In these simulations, all axonal actin monomers and filaments – nucleating and elongating according to the abovementioned model-parameters – were initially considered to be in the “dark state”. Thereafter, a 15 µm axon-segment was computationally “photoactivated”, converting all the actin in this region into an “activated state” (see schematic in **fig. 5A**). Distribution of the photoactivted actin monomer population for an axon is shown in **figure 5B**, left panel. The virtual photoactivated actin population is subsequently tracked as it diffuses along the axon, occasionally translocating when monomers get incorporated into the actin trails (see **Supp. Materials and Methods** for more details). A schematic of the expected anterograde or retrograde intensity center shifts from simulated photoactivation experiments is depicted in **figure 5B**, right.

**Fig 5:**
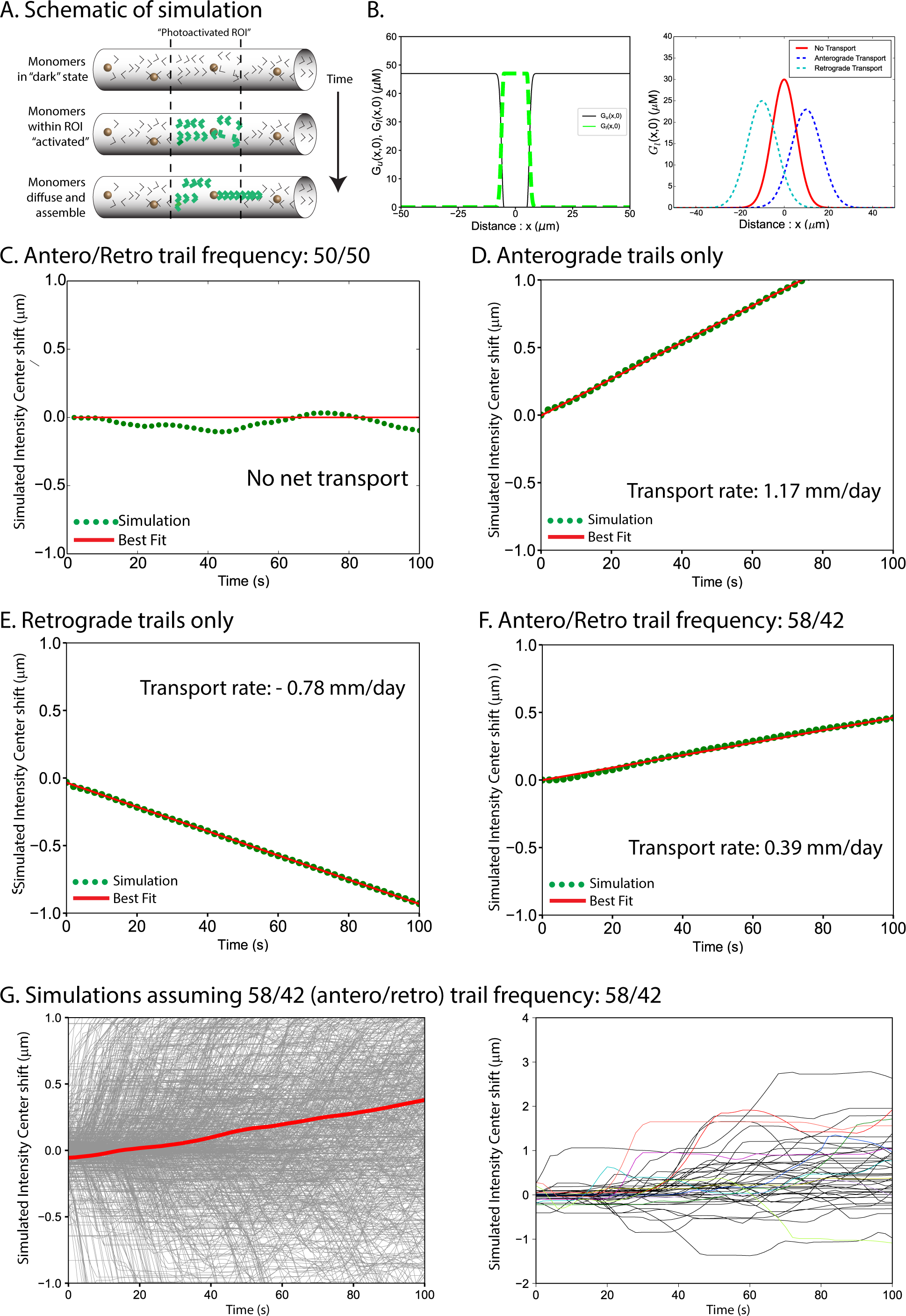
Virtual photoactivation paradigm to evaluate contribution of actin hotspots/trails to slow transport. **(A)** Simulation of pulse chase imaging. Initially all actin is in monomeric state (top panel). After equilibration of the system, axonal actin within a 15 µm region is photo activated (center panel, green arrowheads represent photoactivated actin). Subsequently, the fluorescent actin either diffuses, or translocates after incorporation into trails (bottom panel; see “methods” for more details). **(B)** Left panel shows labeled (green dotted line) and unlabeled (black solid line) actin concentration profiles along the axon shortly after photoactivation. Right panel shows expected outcomes of photoactivation simulations: For no net transport, the center of the labeled G-actin distribution is expected to be at its initial position (red line). For net anterograde or retrograde transport, the center is expected to move anterogradely (blue line) or retrogradely (cyan line), respectively. **(C)** Simulated intensity center shift for equal anterograde and retrograde trail nucleation rates. The center of fluorescent G-actin (green points) fluctuates about its initial position x = 0 indicated by the red line. Results shown are averaged over 100 simulation runs. **(D)** Simulated intensity center shift when only anterograde trails are nucleated. The center of fluorescent G-actin (green points) moves anterogradely at a rate of 0.0135 µm/s (1.17 mm/day). Results averaged over 1000 simulation runs and fitted to a straight line (red line). **(E)** Simulated intensity center shift when only retrograde trails are nucleated. The center of fluorescent G-actin (green points) moves retrogradely at a rate of – 0.009 µm/s (– 0.78 mm/day). Results averaged over 1000 simulation runs and fitted to a straight line (red line). **(F)** Simulated intensity center shift for 58/42% bias of anterograde-retrograde trail frequencies. Center of fluorescent G-actin (green points) moves anterogradely at a rate of 0.0045 µm/s (0.39 mm/day). Results averaged over 1000 simulation runs and fitted to a straight line (red line). **(G)** Individual runs of simulated pulse chase experiment (without averaging). Left panel: 1000 individual simulation runs (grey lines) and the averaged (red line) intensity center shift. Right panel: 50 individual runs of pulse chase experiment.

**Figures 5 C-F** show cumulative average shifts from simulations, where we altered the number of elongating anterograde and retrograde trails (frequency) in axons. As expected, when the number of elongating anterograde and retrograde trails was equal (i.e. a 50/50 frequency), there was no net transport (**fig. 5C**, 100 simulations). If only anterograde trails were allowed, the entire population was transported anterogradely at a rate of 0.0135 µm/s, or 1.17 mm/day (**fig. 5D,** 1000 simulations). If only retrograde trails were allowed, the actin population had an overall retrograde bias at a rate of -0.009 µm/s, or -0.78 mm/day (**fig. 5E,** 1000 simulations). However, an anterograde/retrograde bias of 58/42% – as seen in our imaging data – led to an overall transport rate of 0.0046 µm/s, or 0.39 mm/day (**fig. 5F**, 1000 simulations). Note that the transport rates from these virtual experiments are reminiscent of the data from our real imaging experiments (**fig. 3**, rate = 0.004 µm/s or 0.34 mm/day); and are also comparable to rates determined by pulse-chase radiolabeling of central nervous system neurons in vivo [for example, mean actin transport rates in mouse cortico-spinal neurons was ~ 0.4 mm/day, (Oblinger, 1988)]. Raw intensity-center shifts from all 1000 simulations (the “58/42 ratio” group) are shown in **figure 5G**, left. **Figure 5G**, right shows 50 random intensity center shift patterns from the entire dataset. The stochastic pattern of intensity center shift – seen in both real and virtual datasets (compare **fig. 3C** to **fig. 5G** – right) – is likely due to arbitrary highlighting of anterograde/retrograde trails that happened to be elongating in the photoactivated zone at the time of activation.

### Biased polymerization: An Unusual Mode of Cytoskeletal Transport

The bulk of cytoskeletal proteins are synthesized in the neuronal soma and transported into the axon, where they play important structural and signaling roles. Though the form in which these proteins are transported – monomer or polymer – was heavily debated in the 90’s (Baas and Brown, 1997; Hirokawa et al., 1997); axonal transport of cytoskeletal polymers has now been unequivocally demonstrated (Yan and Brown, 2005). A model has emerged from these experiments, where cytoskeletal monomers are assembled in the neuronal soma, and the assembled polymers are translocated along the axon using the same motors as used by vesicles – kinesins and dyneins (Baas, 2002; Brown, 2000). However, unlike vesicles that move persistently, individual cytoskeletal polymers move much more infrequently and intermittently, making the overall population slow – the “Stop and Go” model of cytoskeletal transport (Brown, 2000). Though there is good evidence that neurofilaments and microtubules are conveyed in axons as polymers, mechanisms underlying the axonal transport of actin have been mysterious for some time [reviewed in (Galbraith and Gallant, 2000)]. One striking difference between actin and other axonal cytoskeletal proteins is that actin is highly dynamic. Neurofilaments are extremely stable polymers with half-lives of months to years (Barry et al., 2007); and neuronal microtubules also quite stable (Baas et al., 1991). In contrast, actin at the axonal growth cone is extremely dynamic (Gallo and Letourneau, 2004), and our data also indicate a similar dynamism in the axon shaft [**fig. 1, 2**; also see (Ganguly et al., 2015)].

Nevertheless, given the precedence of polymer transport in axons, we began these experiments expecting to see actin filaments translocate in axons. However, we were surprised to see unusual on/off actin dynamics, and elongating (not translocating) actin polymers. Collectively, our imaging and modeling data advocate a new mode of cytoskeletal transport, where biased assembly and polymerization can lead to a slow, anterograde movement of the population. Note that this “Biased Polymerization” model is fundamentally distinct from the Stop and Go model, as there is no polymer-sliding. In hindsight, most prevailing models of axonal transport do not take dynamic assembly/disassembly into account. Though this may have minimal consequences for highly stable proteins like neurofilaments, monomer/polymer exchange is likely to have dramatic consequences for the transport for unstable cytoskeletal proteins such as actin. We propose that other dynamic cytoskeletal proteins in neurons (and perhaps also non-neuronal cells) may employ similar strategies for translocation. Polarized assembly of actin is also seen in other contexts, and the precise molecular mechanisms creating such bias are generally unclear (Allard and Mogilner, 2013). The underlying reason for the polarized elongation of actin trails in axons is also unclear, and an inventory of the molecules involved in triggering the hotspots/trails is likely necessary to solve this puzzle.

### MATERIALS AND METHODS

## Neuronal cell cultures, transfection

All mouse procedures were approved by the University of Wisconsin committee on animal care and were in line with NIH guidelines. Hippocampal cultures were obtained from brains of postnatal (P0–P2) CD-1 mice (either sex) and plated on 35 mm MatTek glass bottom dishes as previously described (Ganguly and Roy, 2014). In Brief, MatTek glass bottom dishes were coated with 100µl of 1mg/ml of Poly-D-lysine in 0.1M borate buffer for 2 hrs at room temperature, washed thrice with ddH_2_O, and air dried. Hippocampi from P0-P1 mice were dissected in ice-cold dissection buffer (HBSS, 4.44 mM D-glucose, and 6.98 mM HEPES) and incubated in 0.25% Trypsin-EDTA for 15 mins at 37°C water bath. After blocking in 30% FBS/1X PBS, neurons were resuspended in Neurobasal/B27 media with 10%serum, dissociated, and plated at a density of 25,000 neurons/100µl of plating medium. Neurons were allowed to mature for 7-9 days in Neurobasal/B27 media (supplemented with 2% B27 and 1% GlutaMAX) in an incubator at 37°C and 5% CO_2_. Neurons were transfected with indicated fluorescent proteins with Lipofectamine 2000 (Thermo Fisher Scientific). For Utr-CH constructs, 0.3 µg of DNA was used while for all other constructs 1.2 µg of DNA was used. 12-16 hrs after transfection, neurons were transferred to an “imaging buffer” (Hibernate-E-Low Fluorescence media; Brainbits, LLC, supplemented with 2% B27, 2 mM GlutaMAX, 0.4% D-glucose, and 37.5 mM NaCl), and imaged at 35.5–37°C (on a heated stage chamber, model STEV; World Precision Instrument, Inc.)

### Imaging and Image analyses

Live-imaging experiments were performed on an inverted epifluorescence microscope (Eclipse Ti-E; Nikon) equipped with CFI S Fluor VC 40× oil (NA 1.30; Nikon) and CFI Plan Apochromat VC 100× oil (NA 1.40; Nikon) objectives. An electron-multiplying charge-coupled device camera (QuantEM:512SC; Photometrics) and LED illuminator (SPECTRA X; Lumencor) were used for most experiments. For imaging axonal actin, low GFP:Utr-CH expressing neurons were selected using specific criteria, as described in (Ganguly et al., 2015; Ladt et al., 2016). Axons were identified by morphology, and only neurons with unambiguously identified axons were selected for imaging (Ganguly and Roy, 2014; Roy et al. 2011). Axonal actin was imaged at 20% LED power (400ms exposure), at the rate of one frame per second for a total duration of ten mins. For near simultaneous dual-color imaging, exciting LED lights were rapidly switched (within microseconds) using the SPECTRA X LED illuminator. A dual emission filter cube (Chroma Technology Corp.) was used to collect GFP/RFP emission with subpixel registration. Details of the photoactivation set up has been previously described (Roy et al. 2011). Briefly, a photoactivation region of interest was selected along axons, the PAGFP:Utr-CH was photoactivated for 1 s using the violet LED, and the GFP fluorescence was imaged at two frames per second (with the 100× oil objective). Kymographs were generated using the Kymograph function in the MetaMorph software (Molecular Devices, Sunnyvale, CA). Methods used for quantification of actin hotspots and trails have been previously described in detail (Ganguly et al., 2015). The intensity-center assay was performed using algorithms written in MATLAB [MathWorks; see (Roy et al., 2011; Scott et al., 2011)]. Briefly, after photoactivation, the videos were background-corrected, and the photoactivated ROI was cropped. Intensity line scans along the axon were generated for each frame in the video, and the maximum intensity point (intensity center) was calculated. All data were plotted in Prism for display.

### Actin barbed end labeling in microfluidics

Neurons were grown in a microfluidics device RD450 (Xona microfluidics) as per manufacturer’s instructions with a few modifications. New microfluidic devices were washed with 1% Alconox, followed by thorough washing with ddH_2_0. Thereafter, devices were incubated with 70% ethanol for 30 minutes, washed with sterile distilled water and air dried. The devices were then UV sterilized for 15 minutes. Meanwhile, 60 mm glass bottom MattTek dishes with 20 mm microwell diameter were coated with 1ug/mL Poly-D-Lysine and 10ug/uL of Laminin (Sigma) for 2 hours, and then washed and air dried. UV sterilized microfluidic devices were kept in the microwell of the dish. Before neurons were plated on the device, both main chambers of the device were equilibrated with Neurobasal/B27 media (supplemented with 2% B27 and 1% GlutaMAX) for five mins. Subsequently, Neurobasal medium was removed and approx. 400,000 hippocampal neuronal cells plated on the one side of the chamber (somatodendritic chamber). By days in vitro (DIV) 3-4, axons were found to enter into the other chamber (axonal chamber).

The barbed end labeling protocol was adapted from (Marsick and Letourneau, 2011). First, 0.45 µm rhodamine labelled actin (APHR, Cytoskeleton) solution was prepared in permeabilization buffer (138 mM KCl, 10 mM PIPES, 3 mM EGTA, 4 mM MgCl_2_, and 1% BSA, pH = 6.9) containing 0.025% saponin and 0.2mM ATP. For rhodamine-labeling experiments using microfluidics devices, 100 µl of rhodamine-actin solution was loaded into the axonal chamber for two mins. Immediately thereafter, both chambers were fixed with 4% paraformaldehyde, 0.05% glutaraldehyde and 10% sucrose made in 1X PBS for five mins and washed thrice. The Microfluidic device was then carefully removed without damaging the neuronal processes. Neurons were then permeabilized with 0.2% Triton X in PBS, washed and incubated 1µM phalloidin (Life technologies) for an hour. For rhodamine-labeling experiments using CAD cells, the cells were grown at a density of 30,000 cells/100 µl in DMEM/F12 (Thermo Fisher Scientific) with 8% FBS and 1% Pen-Strep (Thermo Fisher Scientific). One day after plating, cells were incubated with rhodamine-actin for two mins followed by fixation, permeabilization with 0.2% Triton X, and phalloidin staining as described above.

### 3D STORM imaging

Rat hippocampal neurons from E18 embryos were cultured on 18 mm coverslips at a density of 6,000/cm2 following guidelines established by the European Animal Care and Use Committee (86/609/CEE) and approval of the local ethics committee (agreement D13-055-8). After 3 to 8 DIV, neurons were fixed, processed and imaged as previously described (Ganguly et al., 2015; Xu et al. 2013). After a brief extraction with 0.25% Triton X-100 and 0.3% glutaraldehyde in a cytoskeleton-preserving buffer, neurons were fixed with 2% glutaraldehyde in the same buffer for 15 minutes before quenching with 0.1% NaBH4 for 7 minutes, blocked and stained with primary and secondary antibodies in “immunocytochemistry buffer” (0.22% gelatin, 0.1% Triton X-100 in phosphate buffer), then incubated overnight at 4°C with phalloidin-Alexa Fluor 647 (0.5μM in phosphate buffer, Life Technologies). Coverslips were placed in STORM buffer (Tris 50 mM pH 8, NaCl 10 mM, 10% glucose, 100 mM MEA, 3.5 U/mL pyranose oxidase, 40 μg/mL catalase) and imaged on a N-STORM microscope (Nikon Instruments). Phalloidin (0.1-0.25 μM) was added in the STORM medium to mitigate progressive unbinding from actin filaments during imaging ("phalloidin-PAINT"). A series of 60,000 images (67 Hz frame rate) was acquired at full power of the 647 nm laser, with progressive reactivation with the 405 nm laser. Sequences of images were processed for localizations using the N-STORM software and 2D-projections of the 3D-STORM data was generated using the ThunderSTORM plugin for ImageJ (Ovesny et al., 2014).

### Modeling

To model the dynamics of actin trails growing in the axon, we incorporate the available experimental data of their nucleation rates, elongation velocities and length statistics in a 1-d mathematical model. In our model, hotspots are uniformly spaced at a distance of 3.6 *μm* and located linearly along the axon (**fig. 4A**, top panel). Using the data from the PAGFP:Utr-CH imaging experiments, the anterograde and retrograde trail nucleation rate of each hotspot is calculated as *r*_*n,a*_ = 0.001885/*s* and *r*_*n,r*_ = 0.001381/*s*, respectively. After the formation of a stable actin nucleus, the trail grows by incorporating monomers from the available pool of G-actin in the axon (**fig. 4D**, bottom panel). The nucleation and subsequent growth of actin trails are modeled by Markov processes. Probabilities of trail nucleation and of competing association and dissociation reactions are calculated at each time step using the nucleation rates *r*_*n,a*_ and *r*_*n,r*_ and the established reaction rates summarized in **figure 4A** (Pollard and Borisy, 2003). The effect of incorporating G-actin from the axonal pool into trails and the release of actin back into it, is modeled as a one-dimensional diffusion equation with sinks and sources and periodic boundary conditions. We use a diffusion coefficient of G-actin monomers in the cytoplasm of 6 *μm*^2^/*s* (McGrath et al., 1998) and a basal G-actin concentration of 47 *µ*M (see **Results** section). The trail collapse mechanism is modeled as an instantaneous process where once an actin trail reaches a pre-set length, it collapses and deposits its monomer content in the axonal pool of G-actin. We discretize the axon into segments of length ∆*x*(~ 0.027 *μm*) and the time into intervals of 10^−5^*s* and solve the diffusion equation by using a finite-difference method (Crank, 1979) using custom-made algorithms compiled using the commercially licensed Intel^®^ Fortran Compiler.

To quantify the overall rate of axonal actin transport, we computationally simulate a fluorescence pulse chase imaging experiment. In this experiment, all actin (globular and filamentous) present within a central region of interest (ROI) of length ~ 15 *μm* (corresponding to the experiments) is photoactivated. The fluorescently activated actin population is subsequently tracked as it diffuses in the axon; occasionally translocating when incorporated into trails. We simulate actin trails in an axon of length *L* = 1000 *μm* and radius *r* = 85 *nm*. At the start of the simulation (*t* = −50*s*), actin is present only in monomeric form. The system equilibrates over time by nucleating trails from hotspots reaching a steady state. At *t*=0 *s* (i.e. 50 seconds after the start of the simulation), both G-actin and F-actin is photoactivated in a central zone of the axon (Fig. 5B). This zone is chosen to be about 15*µm* in length, in accordance with the experimental procedure. The length of the axon is chosen to be much larger than the length of the activation window, so that the activated population of actin can be tracked for a relatively long period of time (~ 100 *s*) before approaching the ends of the axon. The subsequent spatiotemporal dynamics of both the fluorescently active and the inactive G-actin is modeled for each species separately by a one-dimensional diffusion equation in conjunction with the stochastic elongation dynamics of the trails. The two species, however, interact as they both are incorporated into actin trails and transported. We track the population of fluorescently active actin over time as its distribution broadens diffusively and the center is translocated along the long-axis. To quantify the transport rate, we calculate the velocity of the fluorescence center (see **Supplemental Materials and Methods**) by averaging over hundreds of independently seeded simulation-runs.

